# SFPQ Promotes Homologous Recombination via mRNA Stabilization of RAD51 and Its Paralogs

**DOI:** 10.1101/2025.09.08.674956

**Authors:** Sofia Gotthold, Keile R. Hansen, Andrew N. Brown, Soham P. Chowdhury, Hannah I. Ghasemi, Amanda C. Yoon, Christine M. Joyce, Julien Bacal, Brooke M. Gardner, Chris D. Richardson

**Author notes:** Corresponding Author: Chris D. Richardson Molecular, Cellular, and Developmental Biology, University of California, Santa Barbara, 1115 Life Sciences Building, Santa Barbara, CA 93106-9625.

## Abstract

Double-strand break (DSB) repair occurs through non-homologous end joining (NHEJ) or homologous recombination (HR). To identify non-canonical factors that influence DSB repair outcomes, we parsed data from pooled genetic screens. Through this approach, we identified the splicing factor SFPQ, which has been previously reported to associate with DSBs and promote repair. Here, we show that SFPQ depletion alters DSB repair via HR. However, in contrast to other published work, we find that SFPQ does not localize to DSBs but instead stabilizes the expression of RAD51 and its paralogs independently of p53 activation or DNA damage. Our findings suggest that SFPQ contributes to constitutive DSB repair by maintaining RAD51 paralog mRNA stability rather than through direct interaction with DSBs or RAD51 protein and highlight indirect mechanisms by which RNA-binding proteins can influence genome stability.

## Introduction

The ability of cells to preserve genome integrity is fundamental to cellular homeostasis. Failure to properly repair DNA damage can lead to chromosomal instability, mutations, cell death, or inappropriate growth [1–4]. Among the most deleterious forms of DNA lesions are DNA double-strand breaks (DSBs) which arise from both exogenous sources, such as ionizing radiation, and endogenous processes like transcription, replication stress, and oxidative metabolism [5,6]. Because unrepaired or misrepaired DSBs can cause disease or death, cells have evolved robust DSB repair mechanisms, which cluster into non-homologous end joining (NHEJ) and homologous recombination (HR) repair pathways. DSB repair pathway choice is tightly regulated: in response to a DSB, repair is channeled toward either NHEJ or HR [1,7]. NHEJ repairs DSBs by ligating the two broken ends of the DNA molecule together, which can lead to insertion or deletion of sequence, or chromosomal translocations [8]. This repair process is predominant during the G1 phase of the cell cycle. In contrast, HR uses a sister chromatid –an exogenous homologous DNA molecule– as a template to faithfully restore the broken DNA sequence, which limits it to late S or G2 [1,7,9].

Key decision points between these two pathways are likely influenced by 5’ DSB end resection, which permits the loading of RAD51 onto single-stranded DNA (ssDNA) exposed by resection. These processes convert double stranded DNA (dsDNA) at a DSB into a ssDNA nucleoprotein filament that facilitates homology search and strand invasion during HR [4,6,10,11]. Resection and RAD51 loading also prevent NHEJ because the ligase, LIG4, requires double-stranded DNA as a template. The core events in NHEJ and HR have largely been characterized *in vitro*. However, screens for factors required for DSB repair can identify players that do not have a known catalytic role in these processes. Some of these uncharacterized factors likely have *bona fide* roles in DSB repair. For example, the RAD51 paralogs, RAD51B, RAD51C, RAD51D, XRCC2, and XRCC3 act to promote the assembly and stabilization of the RAD51 filament [12]. In living organisms, the BCDX2 complex (which includes RAD51B, RAD51C, RAD51D, and XRCC2) is essential for repairing DSBs. If any single component of this complex is lost, cells become highly susceptible to agents that damage DNA and show a marked decrease in the formation of RAD51 foci following DNA damage [13]. Additionally, mutations in the genes encoding RAD51B, RAD51C, and RAD51D have been found to be associated with ovarian, prostate and breast cancers [13–15].

In this study, we identify the multifunctional RNA-binding protein Splicing Factor Proline and Glutamine Rich (SFPQ), as a regulator of DSB repair via HR. SFPQ, also known as PSF, is implicated in diverse aspects of RNA metabolism including splicing, transcriptional regulation, R-loop resolution, and RNA transport [16,17]. SFPQ is a key component of paraspeckles; subnuclear condensates that regulate stress responses in viral infection, cancer, development, and neurodegeneration [18]. Importantly, SFPQ dysfunction has been linked to multiple cancers including ovarian, breast, and prostate malignancies; as well as neurodegenerative diseases, such as ALS and frontotemporal dementia [16,19–23]. SFPQ has been previously linked to DSB repair through reported interactions with RAD51 *in vitro* [24] and colocalization with DNA damage-induced foci by laser microirradiation [25,26]. SFPQ has also been implicated in strand invasion during HR, reconstituted *in vitro* [27]. Here, we report that SFPQ depletion significantly impairs recombination, mimicking the effect of RAD51 knockdown. This recombination phenotype for SFPQ has already been reported by others [28], however, contrary to previous reports, we observe that SFPQ does not localize to DSBs. Instead, our data indicate that SFPQ depletion reduces the expression of RAD51 and its paralogs.

Moreover, this effect occurs independently of p53 activation and does not require DNA damage induction. Overall, we propose that SFPQ promotes HR indirectly by binding to and stabilizing RAD51-family mRNAs. Our findings underscore the importance of post-transcriptional regulation in DSB repair and highlight a previously uncharacterized role for SFPQ in maintaining genome integrity via stabilization of mRNAs encoding HR factors.

## Results

### SFPQ depletion influences gene editing outcomes

We reanalyzed data from pooled CRISPRi screens to identify factors that regulate HR [29,30]. SFPQ was genetically required for HR in pooled screens using single- or double-stranded template DNA. Moreover, SFPQ was a top hit in a screen designed to identify genes required to survive nuclease-induced DSBs **(Fig. 1A and 1B)**. Previous publications have found that SFPQ localized to sites of DSB damage and enabled HR, although the exact mechanism by which this happens remains to be understood [28,31,32]. To further test the role of SFPQ or its binding partner, NONO, in HR, we performed a DR-GFP reporter recombination assay. The DR-GFP reporter contains two mutated GFP genes in direct repeats: the upstream copy harbors an I-SceI site, and the downstream copy is truncated. Introduction of Cas9 targeting the I-SceI site induces a DSB in the upstream gene, and repair by HR restores functional GFP, allowing GFP⁺ cells to be quantified by flow cytometry [33]. Using this approach, we observed that depletion of SFPQ reduced HR ∼2-fold and depletion of NONO reduced HR ∼1.6-fold relative to mock treated controls. This was similar to the ∼2-fold reduction in HR seen in the context of RAD51 depletion **(Fig. 1C)**. Together, these findings demonstrate that SFPQ and its partner NONO contribute to HR. To understand how SFPQ regulates the DNA damage response, we next examined its spatial distribution relative to DSBs.

**Figure 1:**
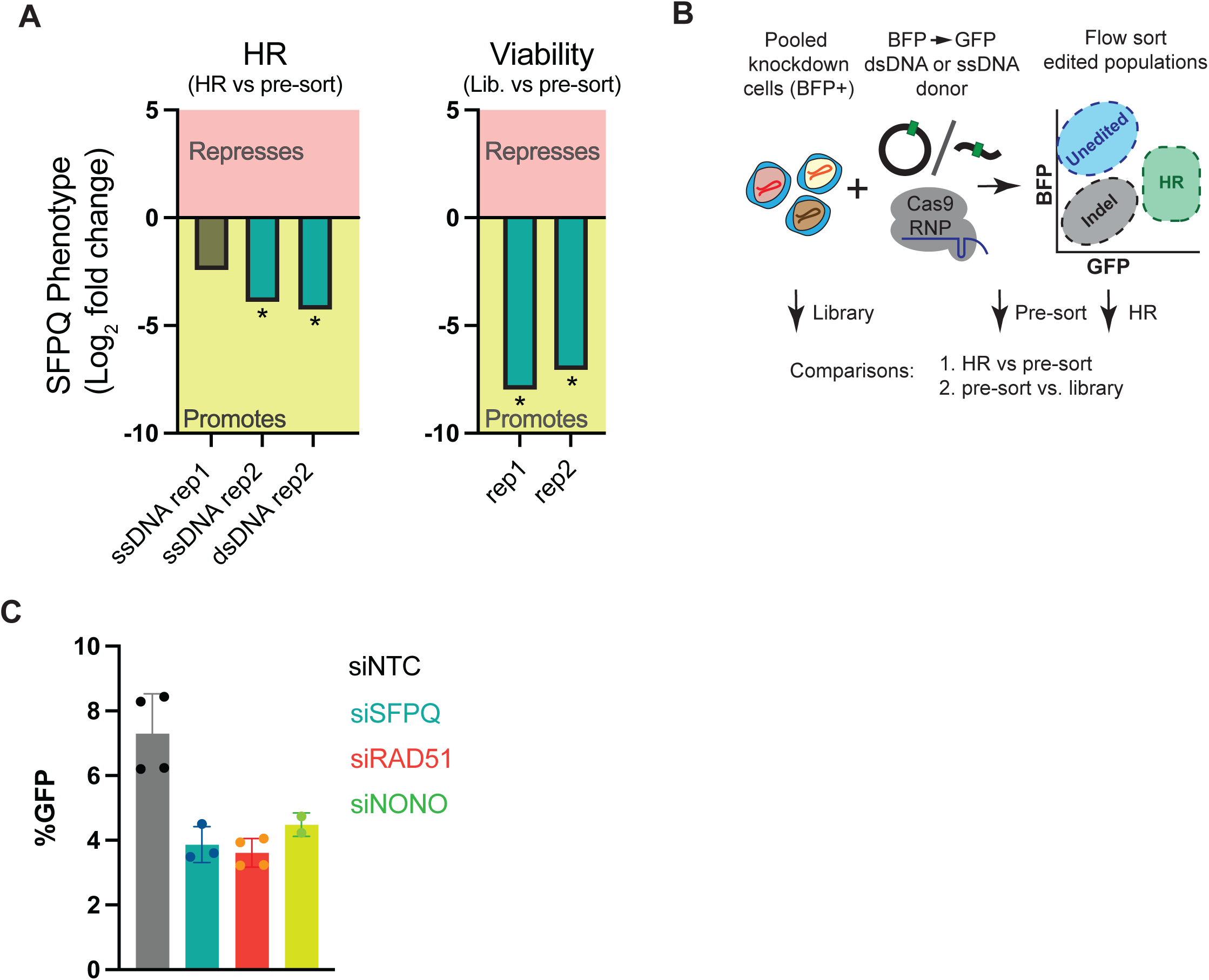
SFPQ depletion influences gene editing outcomes. (A) Column plots showing phenotype scores from pooled genetic screens measuring contribution of genes to HR (left) or viability following DSB induction (right). Data are presented as the log2-fold change in normalized abundance of SFPQ-depleted cells in the experimental (i.e. HR in HR screen, pre-sort in viability screen) relative to control condition. Negative phenotype scores indicate that the gene functions to promote the outcome selected in the screen. * - Mann-Whitney p<0.05. (B) Schematic overview of the experimental workflow used to generate the data shown in panel A. Pooled knockdown cells expressing BFP (BFP⁺) were co-electroporated with a Cas9 RNP complex and either a single-stranded (ssDNA) or double-stranded DNA (dsDNA) donor template converting BFP to GFP. Following editing, cells were subjected to flow cytometry to separate unedited (BFP⁺), indel-containing, and HDR-edited (GFP⁺) populations. Cell populations without DSB induction (library), after DSB induction (pre-sort), or that underwent HR (HR) were processed to recover gRNA abundance and bioinformatically compared to assess depletion or enrichment of gRNAs in specific populations. (C) Percentage of GFP⁺ cells in DR-GFP reporter assays in cells pre-treated with the indicated siRNAs. Bars represent mean ± s.d. from >2 independent biological replicates.

### SFPQ does not colocalize with DSB markers but associates with RNU gene promoters in a break dependent manner

Previous studies report that SFPQ colocalizes to DSBs caused by laser irradiation [26,27]. We therefore measured colocalization of SFPQ with DSB-binding proteins at nuclease-induced DSBs in DIvA (DSB Inducible via AsiSI) U2OS cells. Addition of 4-hydroxytamoxifen (4-OHT) to DIvA cells causes nuclear relocalization of the AsiSI nuclease and induces ∼120 sequence-specific DSBs across the genome [34]. To ensure that we were measuring recruitment of SFPQ to DSBs and not to lesions caused by replication or DSB-replication collisions, we used quantitative image-based cytometry (QIBC) to restrict our colocalization analysis to G2 cells. We treated DIvA U2OS cells with 4-OHT for 24 hours with EdU present for the last 30 minutes of the induction, then immunostained the cells for SFPQ and phosphorylated ATM (a DSB marker [35]), and stained nuclei with DAPI. EdU and DAPI signal were used to determine cell cycle phase, while SFPQ and pATM stains were used to monitor colocalization of factors at DSBs **(Fig. 2A)**. In G2 nuclei, pATM strongly colocalized with DSBs (defined by γH2AX, correlation coefficient R=0.78). In contrast, we observed negligible SFPQ-pATM colocalization in the absence of breaks (R=0.57), and the correlation coefficient for pATM and SFPQ in fact decreased with DSB induction (R=0.4) (**Fig. 2B)**. SFPQ intensity and foci number did increase following DSB induction (**Fig. S1A**), but these increases did not occur at DSBs and may therefore reflect binding of SFPQ to non-DSB regions of the genome, or RNA-binding events caused by DSBs or 4-OHT.

**Figure 2:**
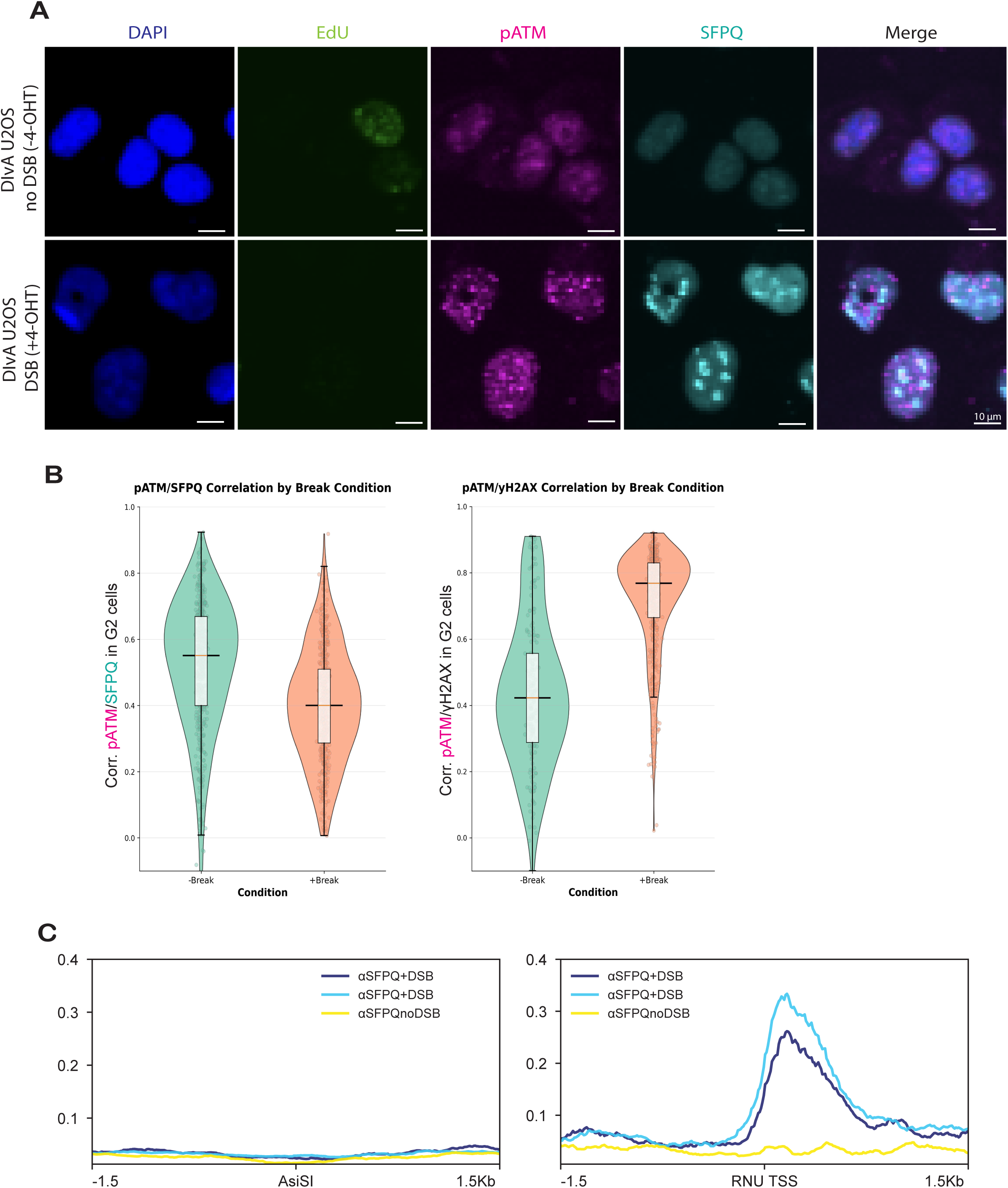
SFPQ does not localize to Double Strand Breaks. (A) Representative images show DAPI (blue), EdU (green), pATM (magenta), SFPQ (cyan), and merged (right) staining in siNTC-treated DIvA U2OS cells under break and no break conditions. Breaks were induced with 4-hydroxytamoxifen for 24 hours, and images were captured using a 10X magnification. Cells (∼20,000 per well) were imaged across three wells per condition (16 fields per well; 48 images total) and quantified in Cell Profiler for nuclear intensity, foci count, and cell-cycle stage based on EdU/DAPI. Data are representative of n=48 images. (B) Quantification of SFPQ–pATM and pATM–γH2AX co-localization in G2-phase cells. Violin plots show correlation coefficients of SFPQ and pATM (left) and pATM and γH2AX (right) in G2 cells with or without DNA breaks. (C) ChIP-seq data representing SFPQ-bound chromatin at 122 defined AsiSI sites under uncut (noDSB) and cut (+4OHT, 4 hours) conditions (left). ChIP-seq data representing SFPQ bound to RNU sites (right). Immunoprecipitation was performed using SFPQ polyclonal antibody. Normalized ChIP-seq signal was plotted for ±1.5 kb around AsiSI sites. SFPQ occupancy profiles are shown for two independent replicates with DSB induction (dark blue and light blue) and for the noDSB control (yellow).

We next performed chromatin immunoprecipitation followed by sequencing (ChIP-seq) to directly measure SFPQ binding to DNA at AsiSI-cut sites 4 hours after DSB induction. Consistent with our microscopy results, we detected no enrichment of SFPQ at AsiSI-induced DSBs. Unexpectedly, we found that SFPQ binding increased at genomic sites encoding small nuclear RNAs (RNUs), a class of non-coding RNAs involved in splicing [36], following DNA damage compared to no-break controls **(Fig. 2C)**. These results indicate that, despite previous reports, SFPQ does not localize to DSBs, but instead is recruited to RNU genes in a break-dependent manner; with the strongest signal observed at RNU7-1, RNU5E-1, RNU1-6, WDR74 (RNU2-2), RNU1-1, RNU12, RNU1-2, SIRT4, RNU1-3, and RNU5B-1. These findings suggest that SFPQ may support HR indirectly, by regulating the splicing or stability of DNA repair factor transcripts, rather than acting directly at the site of the break or regulating transcription of DNA damage response proteins.

### SFPQ knockdown suppresses RAD51 mRNA expression and RAD51 protein levels independently of p53 and DNA damage

Given SFPQ’s role as a splicing factor and recruitment to RNU elements, we examined the effects of SFPQ depletion on the expression of DNA repair genes. Relative to siNTC controls, SFPQ knockdown significantly decreased the transcript abundance of RAD51, RAD51B, RAD51C, RAD51D, XRCC2, and XRCC3, with log₂ fold changes ranging from -0.5 to -1.5 (p < 0.005) **(Fig. 3A)**. Other HR and DSB-repair genes were also downregulated, but the effect size was greatest for RAD51 paralogs. This RAD51 paralog transcript reduction is a steady-state consequence of SFPQ depletion, as RAD51-paralog expression was not DSB-dependent, nor did SFPQ bind to the upstream/downstream regions surrounding these genes (**Fig. S2A-B**). Differential transcript utilization analysis revealed that SFPQ depletion significantly altered the splicing of RAD51B and RAD51C, but not RAD51, RAD51D, XRCC2, or XRCC3 **(Fig. 3B)**. These results indicate that SFPQ promotes HR by sustaining RAD51 family transcript abundance, with additional contributions from regulation of alternative splicing of RAD51B and RAD51C.

**Figure 3:**
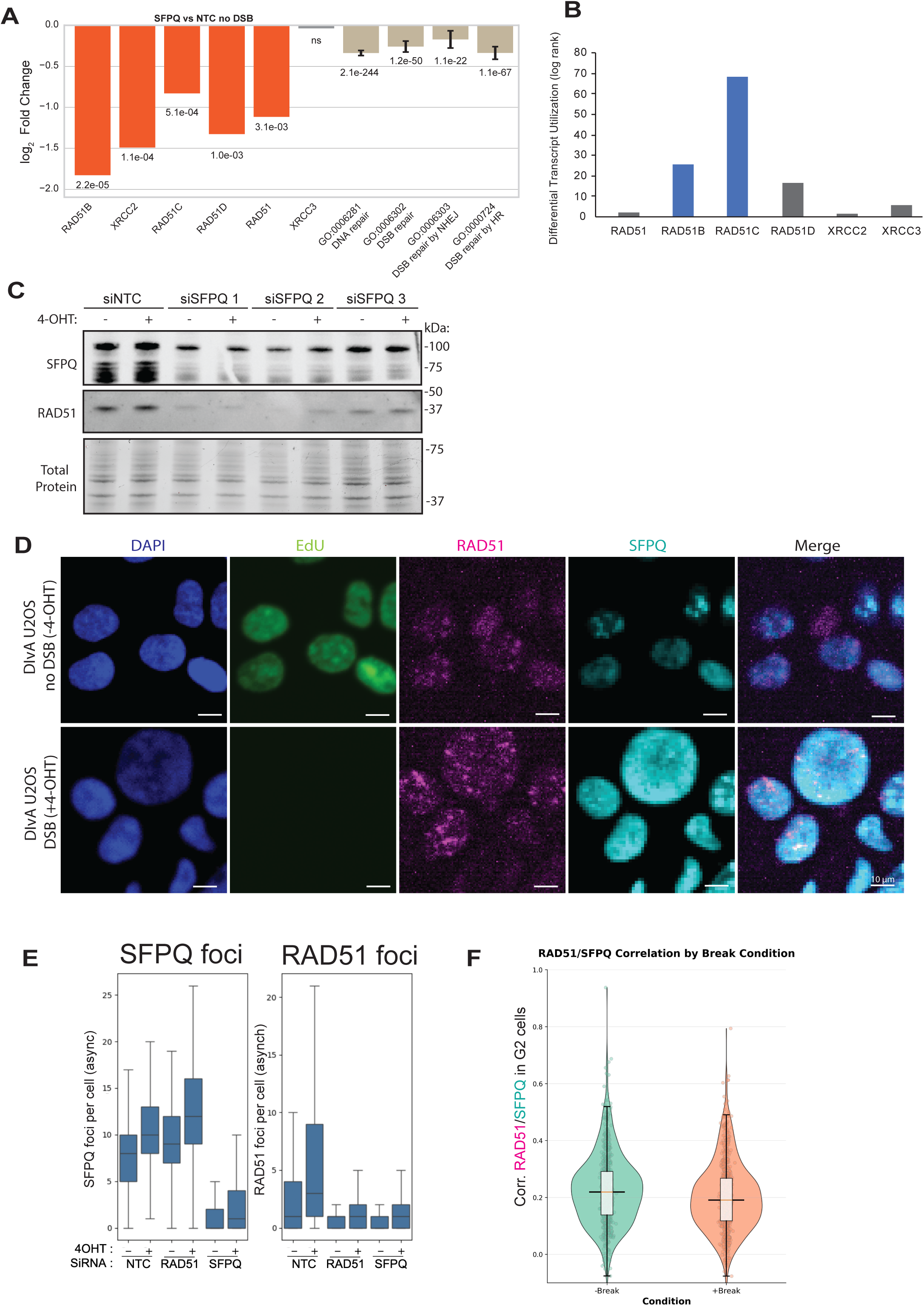
SFPQ knockdown suppresses RAD51 and paralog mRNA expression and RAD51 protein levels independently of DNA damage. (A) mRNA-seq log₂ fold changes of RAD51 paralogs and pooled transcripts in the indicated Gene Ontology (GO) categories in DIvA U2OS cells treated with siSFPQ compared to siNTC control for 72 hours in the absence of DSBs. Data represent the mean of three biological replicates. Individual p-values were adjusted for multiple comparisons. Aggregate p-values were combined by Fisher’s method. (B) Differential transcript utilization analysis for RAD51 paralogs. mRNA-seq data from siNTC versus siSFPQ DIvA U2OS cells were analyzed for transcript isoform usage. Bars represent the likelihood ratio statistic for each gene, with blue bars indicating genes showing significant shifts in transcript utilization (RAD51B, RAD51C) upon SFPQ depletion. Grey bars indicate genes without significant changes. (C) Western blot analysis of SFPQ and RAD51 protein levels of the three biological replicates used for mRNA-seq following siNTC or siSFPQ treatment. Total protein staining is shown as a loading control. (D) Representative images show DAPI (blue), EdU (green), RAD51 (magenta), SFPQ (cyan), and merged (right) staining in siNTC-treated DIvA U2OS cells under break and no break conditions, pre-extracted with CSK. Breaks were induced with 4-OHT for 4 hours. Cells (∼20,000 per well) were imaged across four wells per condition (16 fields per well; 64 images total) and quantified in Cell Profiler for nuclear intensity, foci count, and cell-cycle stage based on EdU/DAPI. (E) Quantification of SFPQ and RAD51 foci per cell in DIvA U2OS cells following siRNA treatment and DNA damage induction. Violin plots show the distribution of foci counts across conditions with or without 4-hydroxytamoxifen (4-OHT) treatment and following transfection with non-targeting control (NTC), RAD51-targeting, or SFPQ-targeting siRNAs. Data are representative of n=64 images. (F) Violin plots showing correlation coefficients of SFPQ and RAD51 in G2 cells with or without DNA breaks. Quantification of SFPQ-RAD51 foci co-localization in G2-phase cells was performed using Cell Profiler analysis of single-cell fluorescence signals.

We next asked whether the transcript-level changes we observed were mirrored at the protein level. Western blotting revealed a reduction in RAD51 protein in siSFPQ-treated cells. This reduction was independent of 4-OHT-induced DNA damage, confirming that SFPQ regulates RAD51 levels independently of DSB induction **(Fig. 3C)**. To further support this finding, we analyzed RAD51 expression and localization using QIBC. Upon DSB induction, RAD51 foci increased ∼2-fold in siNTC cells, consistent with prior DIvA U2OS reports [37,38], whereas SFPQ knockdown blunted this response, resulting in fewer RAD51 foci per cell **(Fig. 3D-E)**. We also observed that RAD51 mean fluorescence intensity was significantly reduced in siSFPQ cells compared to siNTC controls, corroborating the observed depletion of overall RAD51 protein. Despite this reduction, colocalization analysis revealed a low correlation coefficient (R = 0.2) between SFPQ and RAD51 signals, suggesting that the two proteins rarely colocalize and that SFPQ does not function directly at RAD51 foci **(Fig. 3F)**. These results further support a model in which SFPQ modulates RAD51 abundance.

Our mRNA-seq data showed upregulation of CDKN1A (p21), a canonical p53 target [39–41], in siSFPQ-treated cells **(Fig. S2C)**, suggesting that SFPQ depletion might chronically activate the p53 pathway. To test if the reduction of RAD51 in the context of SFPQ depletion is p53-dependent, we treated siNTC and siSFPQ transfected cells with the p53 inhibitor PFT-α [42]. Immunoblotting confirmed effective p53 inhibition, as MDM2 and HSP70 expression were reduced. However, RAD51 protein levels remained suppressed in siSFPQ cells despite p53 inhibition, indicating that the loss of RAD51 upon SFPQ depletion occurs independently of p53 **(Fig. S2D)**. We confirmed this result in p53-null K562 cells [43], where SFPQ knockdown reduced RAD51 protein relative to control levels **(Fig. S2E)**. These results indicate that SFPQ regulates RAD51 and paralog expression through a p53-independent mechanism. Together, these results demonstrate that SFPQ regulates abundance of RAD51 and its paralogs through a mechanism independent of both p53 and DNA damage.

### SFPQ binds 5**′**UTRs to stabilize RAD51-family mRNAs

To test whether SFPQ directly influences RAD51 protein stability, we performed a cycloheximide (CHX) and carfilzomib (Carf) chase assay in DIvA U2OS cells transfected with siNTC or siSFPQ. Cells were treated with CHX (to block translation) or CHX + Carf (to block translation and proteasomal degradation) [44–48] and harvested at 0, 2, and 4 hours. Immunoblotting confirmed that RAD51 undergoes proteasomal degradation [49], as levels were more stable with CHX + Carf rather than with CHX alone in siNTC cells. RAD51 degraded at the same rate in siSFPQ treated cells, but at t=0 of CHX treatment, there was less RAD51 protein to begin with when normalized to total protein (siNTC RAD51 a.u=1.01 vs siSFPQ a.u=0.6) **(Fig. 4A and 4B)**. This data and our mRNA-seq results prompted us to test whether SFPQ regulates RAD51 abundance primarily through interactions with its mRNA.

**Figure 4:**
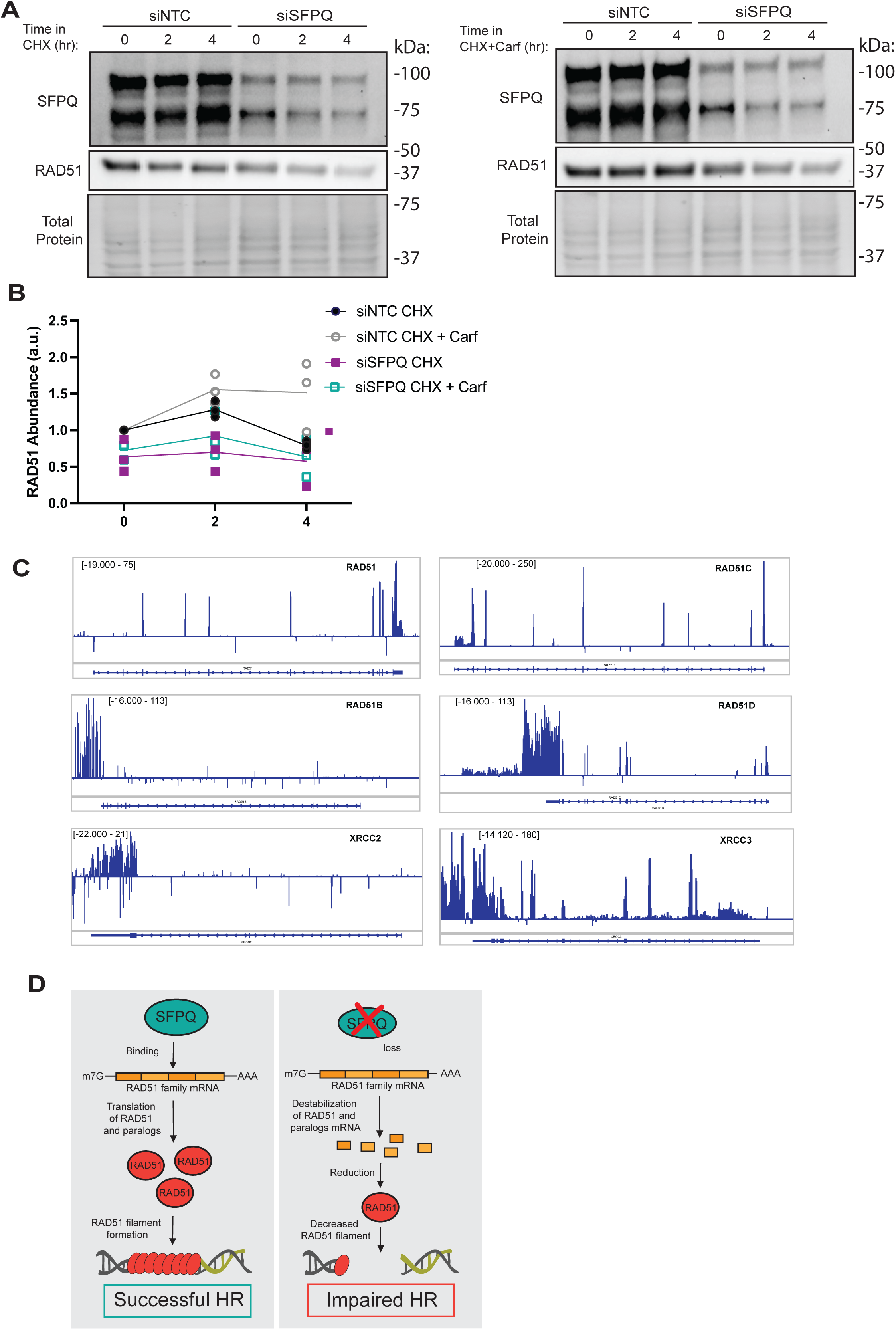
SFPQ binds to exons of RAD51 and paralog transcripts. (A) Cycloheximide (CHX) ± carfilzomib (Carf) protein stability assay in DIvA U2OS cells. Cells were transfected with either non-targeting control (siNTC) or SFPQ-targeting (siSFPQ) siRNAs for 72 h, then treated with CHX alone or CHX + Carf to inhibit protein synthesis and proteasomal degradation, respectively. Lysates were collected at 0-, 2-, and 4-hours post-drug treatment from three independent biological replicates. (B) RAD51 abundance from Fig. 4A normalized to total protein and then to 0 hr. condition. Data points represent individual replicates; lines indicate the mean. (C) RIP-seq analysis of SFPQ binding across RAD51 family paralogs in melanoma cells. Read coverage tracks show SFPQ-associated RNA fragments aligned to the genomic loci of RAD51B, RAD51C, RAD51D, XRCC2, and XRCC3. Peaks indicate regions of enriched SFPQ binding, with annotations of exon–intron structure shown below each track. Model for SFPQ-mediated stabilization of RAD51 mRNA and its impact on homologous recombination (HR). In the presence of SFPQ, the protein binds to RAD51 mRNA, promoting transcript stabilization. Stable RAD51 mRNA ensures sufficient RAD51 protein production, enabling efficient RAD51 filament formation on DNA and supporting robust HR (left). Upon SFPQ loss, RAD51 family mRNAs are destabilized, leading to reduced RAD51 protein abundance. This reduction impairs HR efficiency (right).

We analyzed previously published RNA immunoprecipitation sequencing for SFPQ (RIP-seq) data from melanoma cells [19]. RIP-seq read alignment to the human genome (hg38) revealed strong SFPQ enrichment over exonic, but not intronic regions of RAD51 and its paralogs compared with IgG controls **(Fig. 4C)**. Peak SFPQ-binding occurred in the 5′ untranslated regions (5′ UTRs) of transcripts RAD51, RAD51B, RAD51C, RAD51D and XRCC2 **(Fig. 3A)**. This binding pattern is consistent with previous studies showing that SFPQ regulates transcripts such as CK1α through 5′UTR association [21]. In contrast, XRCC3 shows robust 5′UTR/exonic binding but minimal downregulation, suggesting other transcript-specific buffering mechanisms. Overall, these results support a model in which SFPQ binds to the 5′UTRs of RAD51-family transcripts to promote their stability and/or translation, thereby sustaining RAD51 pathway gene expression and promoting HR.

To test whether this RNA-level coupling is generalizable across cell types, we analyzed RNA-seq from more than 1,500 DepMap cell lines. SFPQ and RAD51 expression were positively associated across tumor lineages (Pearson r = 0.625; p = 6.9×10⁻¹ ), consistent with a model in which SFPQ supports RAD51 expression **(Fig. S4A)**. Together, the DepMap co-expression landscape and our perturbation data support a broad coupling between SFPQ and RAD51 genes.

## Discussion

In this study, we identify the RNA-binding protein SFPQ as a critical regulator of HR through an RNA-centric mechanism that does not involve direct localization of SFPQ to DSBs. Across assays, SFPQ loss lowers RAD51 protein levels and impairs HR to a degree comparable to RAD51 knockdown **(Fig. 1C)**, supporting a model in which SFPQ sustains the expression of RAD51 paralogs to enable recombination. These findings add to a growing body of evidence implicating RNA-binding proteins (RBPs) in the regulation of genome stability. Notably, RBPs such as FUS, hnRNPU, RBMX and CCAR1 have also been shown to modulate the expression and function of DNA repair genes [50,51].

Contrary to previous reports suggesting that SFPQ may localize to DSBs or interact directly with DNA repair factors [17,26,28] our microscopy and ChIP-seq analyses revealed no enrichment of SFPQ at AsiSI-induced DSBs, nor colocalization with DSB repair proteins, nor DNA damage markers. Instead, we observed that SFPQ interacts with the mRNA transcripts of RAD51 and its paralogs, stabilizing them. Depletion of SFPQ therefore reduces RAD51 family gene expression, including RAD51, RAD51B, RAD51C, RAD51D and XRCC2 genes [10,12] **(Fig 3A)**. RAD51 filament formation is stabilized by the BCDX2 components, and in SFPQ knockdown cells, reduced basal levels of RAD51, RAD51B/C/D, and XRCC2 would be expected to delay nucleation, yield fewer foci, and shorten filament lifetime once formed, consistent with the decrease in RAD51 foci we observe [10,52] **(Fig 3C-D)**. We note that SFPQ depletion has a less pronounced effect on XRCC3 expression, but it significantly reduces RAD51C mRNA levels, thereby perturbing the CX3 complex. We therefore hypothesize that SFPQ deficiency may create an imbalance in the relative abundance of the BCDX2 and CX3 complexes, which in turn affects HR outcomes [52].

Despite SFPQ’s canonical role as a splicing factor, we observed no splicing changes in RAD51 itself upon SFPQ depletion; however, RAD51B and RAD51C (both essential for BCDX2 function) underwent differential transcript utilization. These splicing effects may contribute to reduced filament competence, but they do not fully explain the parallel drop in RAD51 mRNA and protein. Thus, we infer that decrease in RAD51 paralog transcript abundance is the primary consequence of HR reduction in SFPQ-deficient cells, with splicing changes of RAD51B and RAD51C acting as a secondary contributor to the HR defect. Consistent with this model, RNAseq in >1500 cell lines show a correlation between RAD51 and SFPQ transcripts **(Fig. S4A)**. Together, population-level in different cancer cell lines, and perturbation data argue for links between SFPQ and RAD51 expression.

Mechanistically, multiple lines of evidence indicate that SFPQ acts on RAD51 mRNA. First, SFPQ does not enrich at AsiSI-induced breaks by QIBC or ChIP-seq, nor does it colocalize with canonical damage markers **(Fig. 2A–C**). Second, SFPQ does not bind at the TSS or TES of RAD51 or its paralogs **(Fig. S2B)**, but it binds exonic/5′UTR regions of RAD51-family transcripts, and SFPQ loss reduces their mRNA abundance without increasing intron retention, consistent with stabilization of mature mRNAs rather than splicing defects in RAD51 **(Fig. 3F**; **Fig 4C)**. Third, protein-stability assays show that RAD51 degradation kinetics are similar with or without SFPQ, but SFPQ-depleted cells start from a lower baseline, indicating that the primary deficit precedes RAD51 protein turnover **(Fig. 4A–B)**. We therefore propose that SFPQ sustains RAD51 paralog expression by stabilizing mature transcripts.

SFPQ likely contributes to DSB repair beyond its role in stabilizing RAD51 paralogs. First, SFPQ-deficient cells have reduced viability after DSB induction compared to mock-treated cells (**Fig. 1A**). Second, nuclear abundance of SFPQ does increase after DSB induction, but this increase does not occur at DSBs (**Fig. 2A**). We favor a model in which transcriptional changes caused by DSB induction or addition of tamoxifen cause localization of SFPQ to RNA transcripts or other RNA structures in the nucleus. This activity broadly supports viability following DNA damage through mechanisms that are independent from direct activities of SFPQ at the DSB itself.

Overall, our findings reveal a non-canonical function for the splicing factor SFPQ in preserving genome integrity through post-transcriptional stabilization of RAD51 and its paralogs. By stabilizing RAD51-family transcripts and preventing their degradation, SFPQ indirectly promotes HR. Loss of SFPQ disrupts this regulation, leading to diminished RAD51 levels and impaired HR **(Fig. 4D)**. This mechanism has important implications for human disease. In cancer, SFPQ is frequently misregulated, overexpressed, or rearranged as part of oncogenic fusion events (e.g., SFPQ–TFE3) [53] and its loss has been associated with altered splicing and genomic instability. Notably, SFPQ dysregulation has been reported in ovarian, breast, and prostate cancers, that also harbor recurrent mutations or altered expression of RAD51 family members, suggesting that disruption of this SFPQ–RAD51 axis may represent a shared vulnerability across these malignancies [16,20,54–57]. This is consistent with the observed correlation between SFPQ and RAD51 expression. Our study suggests that SFPQ depletion may also suppress HR capacity, potentially sensitizing SFPQ-deficient tumors to DNA-damaging agents. Furthermore, SFPQ mislocalization has been observed in neurodegenerative disorders such as Amyotrophic Lateral Sclerosis, Alzheimer’s and Frontotemporal dementia [23,58], where its functional loss could contribute to genome instability and neuronal vulnerability through similar RAD51-dependent mechanisms [59]. Altogether, our results underscore the significance of RNA-binding proteins, like SFPQ, in DNA repair regulation and open new avenues for targeting post-transcriptional networks in disease contexts marked by genomic instability.

## Supplementary Figures

**Figure S1:**
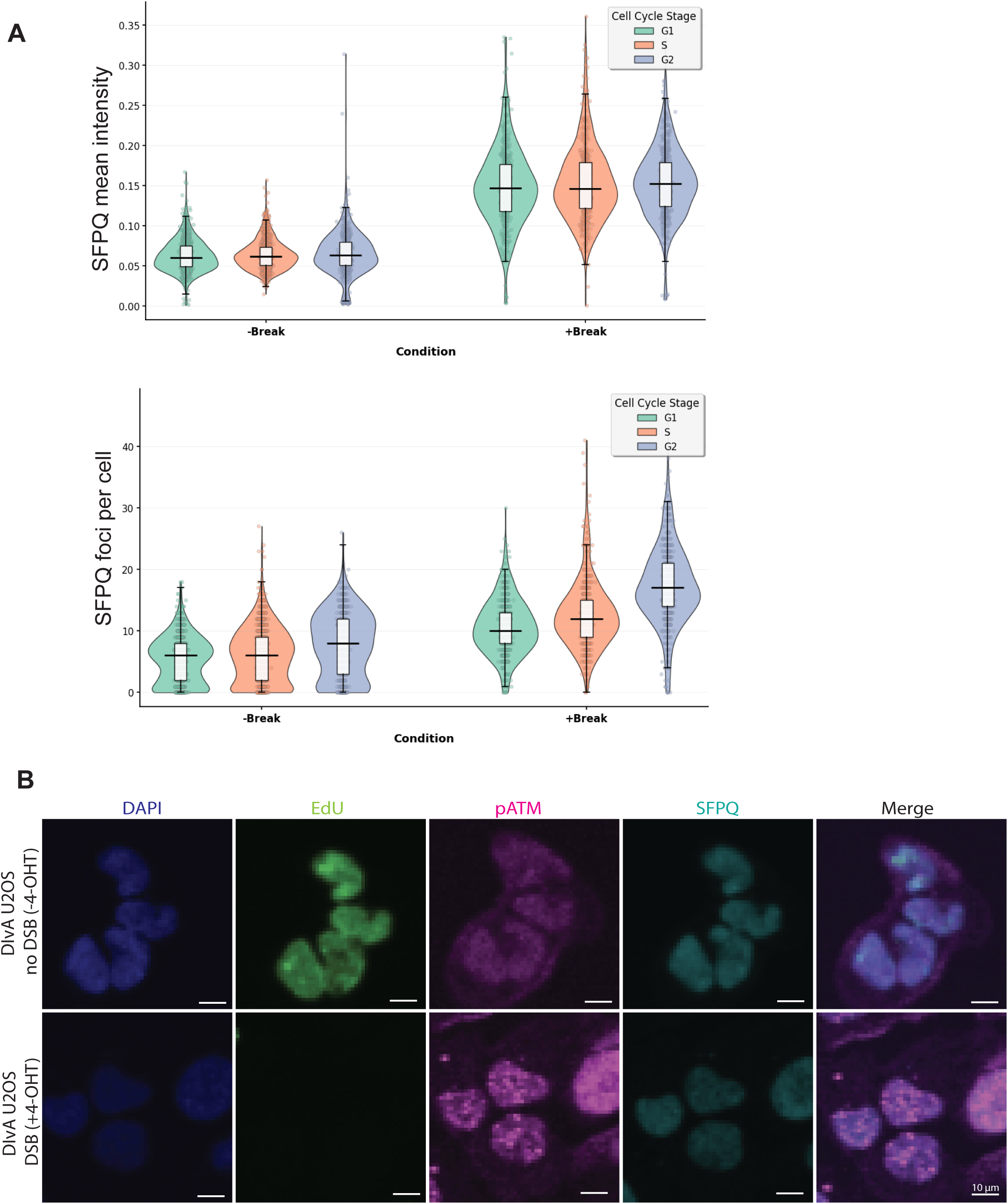
Genotoxic stress elevates SFPQ intensity and foci formation per cell. (A) (Top) SFPQ mean intensity: Violin plots (with embedded boxplots) show the single-cell distribution of nuclear SFPQ mean fluorescence intensity in DIvA U2OS cells under no break (untreated) and break (4-hydroxytamoxifen, 4-OHT) conditions. Each dot is one nucleus; boxplots denote median and interquartile range. Cell-cycle phase (G1, S, G2) was assigned per cell using EdU incorporation (green) and DAPI DNA content (blue). (Bottom) SFPQ foci per cell: Violin plots (with embedded boxplots) show the number of SFPQ nuclear foci per cell under the same conditions and cell-cycle stratification. Quantification: Cells were left untreated or treated with 4-OHT to induce AsiSI-mediated DSBs, then stained for SFPQ (cyan), EdU, and DAPI. Images were analyzed in Cell Profiler to segment nuclei, call SFPQ foci, compute per-nucleus mean intensity and foci counts, and assigned cell-cycle stage from EdU/DAPI features. (B) Non–pre-extracted immunofluorescence staining of pATM and SFPQ in DIvA U2OS cells with or without DSB induction. Cells were left untreated or treated with 4-hydroxytamoxifen (4-OHT) to induce AsiSI-mediated DSBs and stained for DNA (DAPI, blue), EdU incorporation (green), phosphorylated ATM (pATM, magenta), and SFPQ (cyan). Images were acquired without cytoskeletal (CSK) pre-extraction to visualize total nuclear staining patterns. Merged images show nuclear co-localization of pATM and SFPQ signals in the presence and absence of DNA damage.

**Figure S2:**
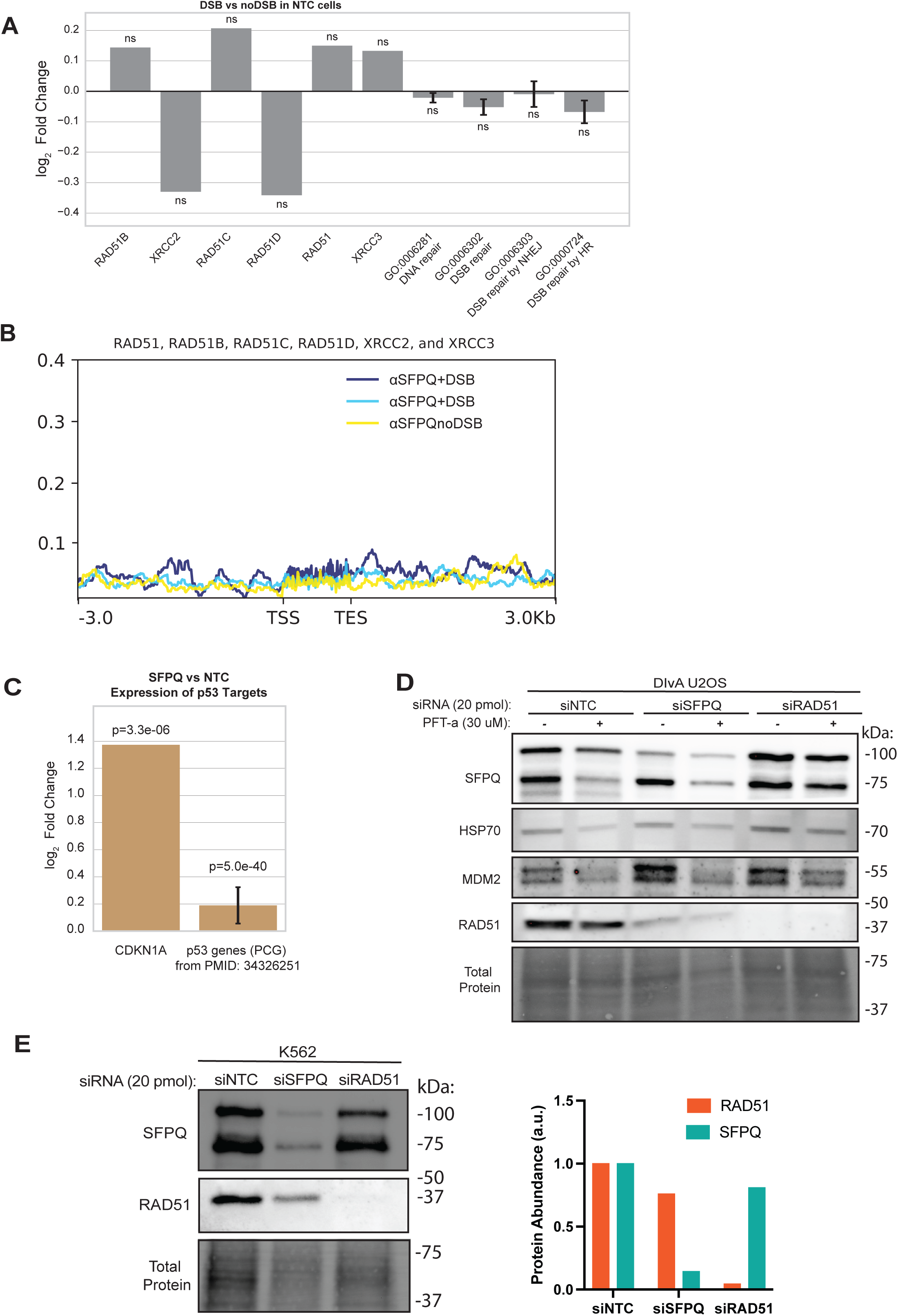
SFPQ knockdown reduces RAD51 expression independently of p53. (A) Differential expression analysis of mRNA-seq data comparing DSB versus no-DSB conditions in siNTC-treated DIvA U2OS cells (n=3 biological replicates). Mean log₂ fold change for the same targets is shown as Fig 3A. No significant expression differences were detected for these targets upon DSB induction in control cells. (B) ChIP-seq data showing SFPQ abundance at sites upstream and downstream of RAD51-paralog genes both without (noDSB) or with (+DSB) 4 hours of DSB induction. Data displayed is the average signal across all 6 RAD51 paralogs. (C) mRNA-seq log₂ fold changes of transcript expression of the indicated gene or GO category in DIvA U2OS cells treated with siSFPQ compared to siNTC control for 72 hours in the absence of DSBs. Data represent the mean of three biological replicates. Individual p-values were adjusted for multiple comparisons. Aggregate p-values were combined by Fisher’s method. (D) Western blot of DIvA U2OS cells treated with siSFPQ with or without p53 inhibition by PFT-α (30 µM) for 24 hours. Lysates were blotted for SFPQ, HSP70, MDM2 and RAD51. (E) (Left) Western blot of p53-null K562 cells treated with siSFPQ. Total protein staining is shown as a loading control. (Right) Quantification of SFPQ and RAD51 normalized band intensities relative to total protein is graphed.

**Figure S3:**
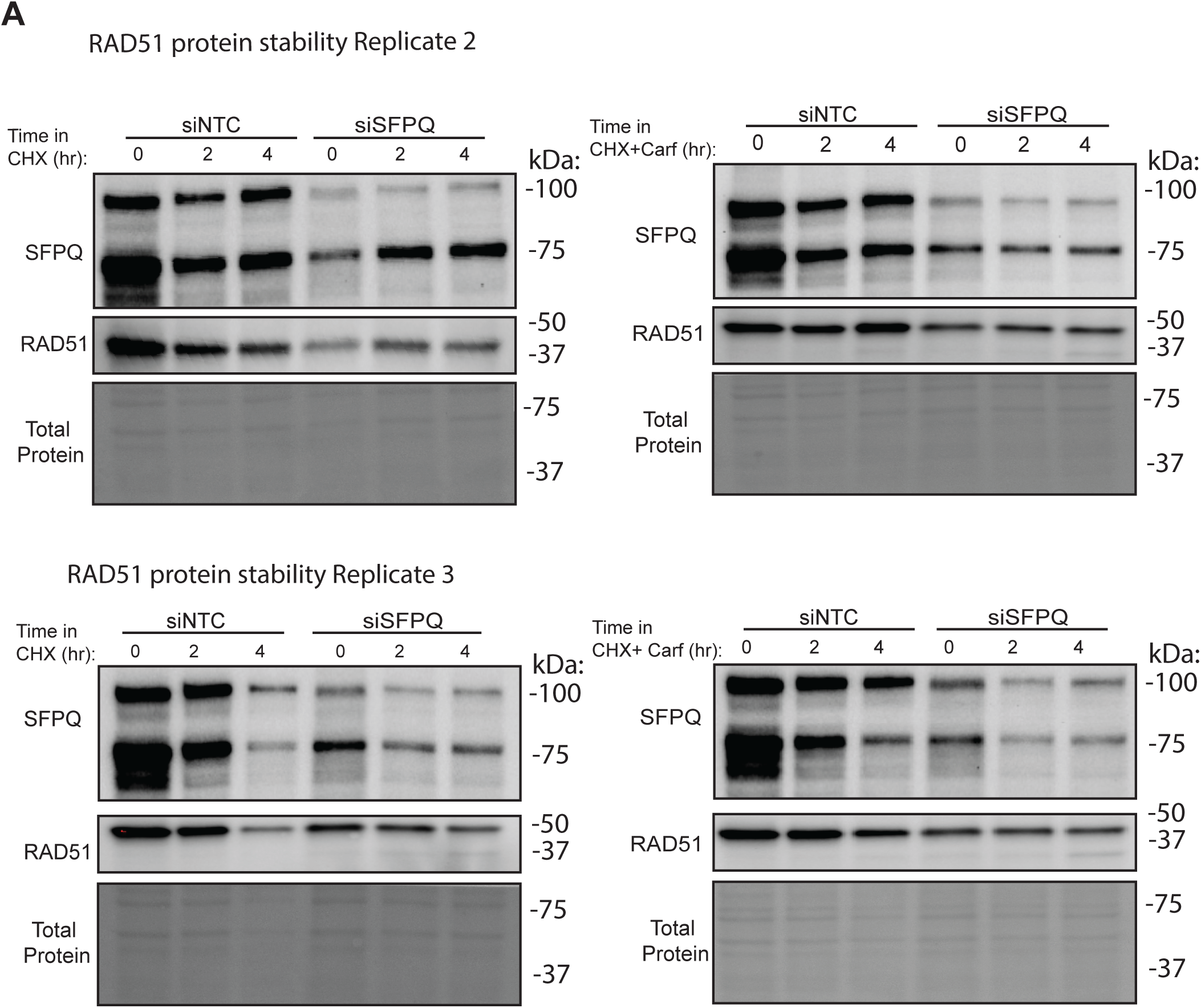
Post-transcriptional regulation of RAD51 by SFPQ. (A) Biological replicates of RAD51 protein stability assays. DIvA U2OS cells were transfected with non-targeting control (siNTC) or SFPQ-targeting (siSFPQ) siRNA for 72 hours, followed by treatment with cycloheximide (CHX) alone or CHX plus carfilzomib (Carf). Lysates were collected at the indicated time points (0, 2, and 4 hours) and RAD51 abundance was measured by Western blot, normalized to total protein.

**Figure S4:**
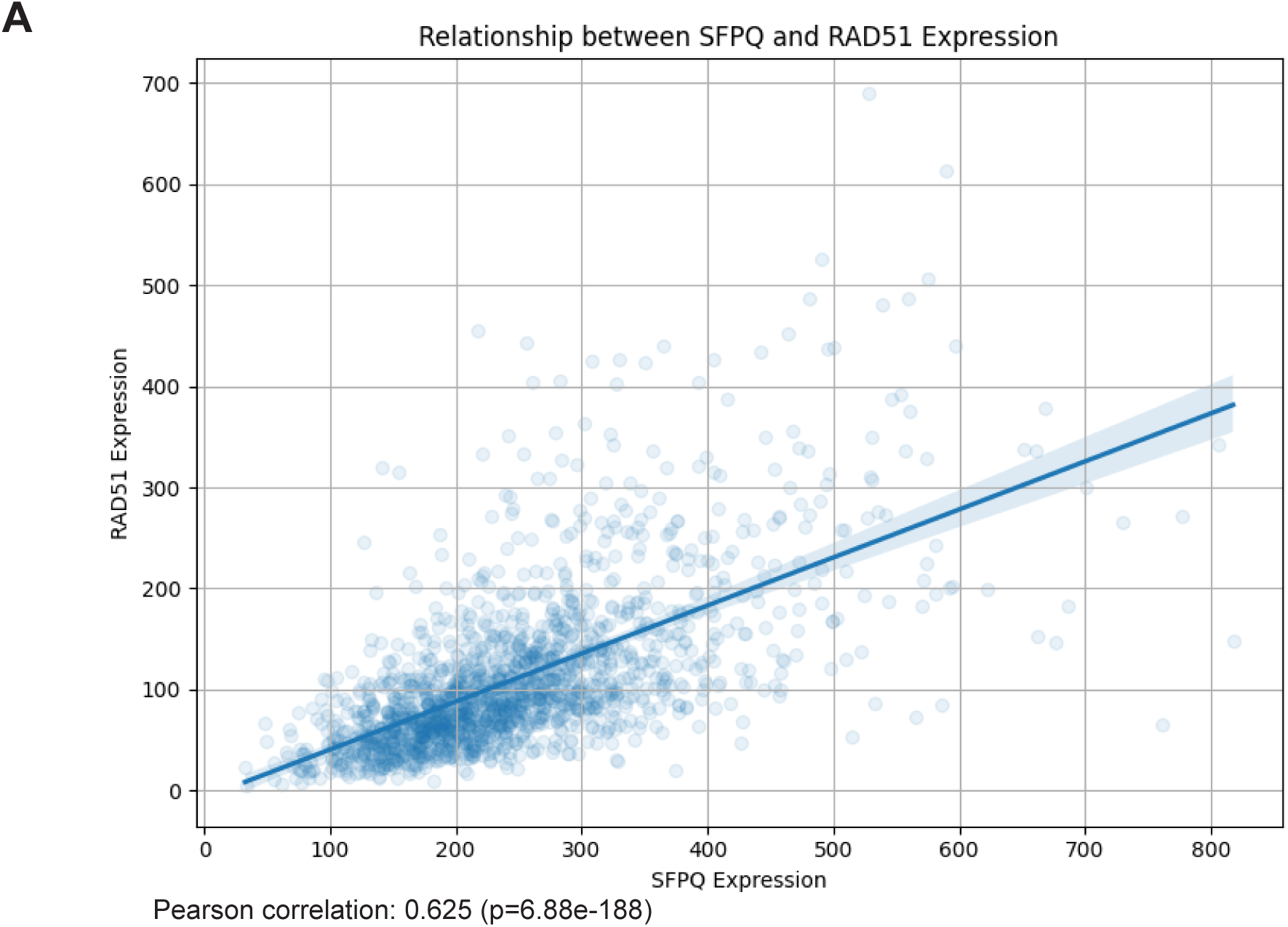
Correlation between SFPQ and RAD51 expression. (A) Scatterplot showing RAD51 expression (TPM) as a function of SFPQ expression across 1684 cell lines available in Depmap. (Pearson r = 0.625; p = 6.9×10⁻¹ ).

## Methods

### Cell lines and culture

DIvA AsiSI-ER-U2OS cells (female; provided by Gaëlle Legube’s laboratory, Center for Integrative Biology, Toulouse, France) were cultured in DMEM GlutaMAX (Gibco, Cat#10569044) supplemented with 10% fetal bovine serum (FBS; R&D Systems, Cat#S11550) and 1% penicillin–streptomycin (P/S; Gibco, Cat#15140122). K-562 cells (female; ATCC) were maintained in RPMI 1640 medium (Gibco, Cat#72400120) containing 10% FBS and 1% P/S. U2OS DR-GFP cells (female; provided by Jeremy Stark’s laboratory, City of Hope, Duarte, California, USA) and wild-type U2OS cells (female; ATCC) were both grown in DMEM supplemented with 10% FBS and 1% P/S. DIvA AsiSI-ER-U2OS and U2OS DR-GFP cell lines were additionally maintained in 1 µg/mL puromycin dihydrochloride (Gibco, Cat#A1113803). All cell cultures were kept at 37 °C in a humidified incubator with 5% CO₂.

### RNA Interference (siRNA)

WT U2OS, DIvA AsiSI-ER-U2OS, and U2OS DR-GFP cells (2–2.5 × 10 per well) were seeded in 6-well plates and transfected 24 h later with 20 pmol siRNA targeting SFPQ, RAD51, NONO, or a non-targeting control (Dharmacon) using Lipofectamine 2000 (Invitrogen, Cat#11668019) in Opti-MEM (Gibco, Cat#31985070) according to the manufacturer’s instructions. Medium was replaced 24 h post-transfection, and cells passaged if necessary. Experimental procedures, including DSB induction, genome editing, and lysate collection, were performed 72 h after transfection.

### Small molecule inhibition

DIvA AsiSI-ER-U2OS cells were treated with small molecule inhibitors, with or without 300 nM 4-hydroxytamoxifen (4-OHT) to induce DSBs. Unless otherwise indicated, treatments were carried out for 4 h prior to ChIP or for 24 h prior to QIBC. For PFT-α (Cat# 506132-5MG), final concentrations of 20 μM or 30 μM were added 24 h before lysate collection.

### Western Blotting

Cells (∼1 × 10 per well) in 6-well plates were washed three times with D-PBS and lysed in 60 μL 2× Laemmli buffer (20% glycerol, 125 mM Tris–HCl pH 6.8, 4% SDS, 0.02% bromophenol blue, 5% 2-mercaptoethanol). Lysates were collected by scraping, sonicated (15 s on/5 s off, total on-time 30 s, 50% amplitude; Branson), boiled at 95°C for 3 min, and centrifuged at 21,100 × g for 5 min. Proteins were resolved on 4–20% SDS-PAGE gels (Bio-Rad, Cat#4568096) and quantified using Stain-Free total protein detection. When necessary, protein loading was normalized by whole-lane densitometry. Proteins were transferred to 0.45 μm LF PVDF membranes via semi-dry transfer (Trans-Blot Turbo, Bio-Rad), blocked for 1 h in 3% BSA in TBS–0.1% Tween-20 (TBS-T), and probed overnight at 4°C with primary antibodies against SFPQ or RAD51 in 3% BSA/TBS-T. Membranes were washed and incubated for 1 h with HRP-conjugated secondary antibodies diluted in 5% NFDM, then visualized using either Pierce ECL 2 (Thermo Scientific, Cat#PI80196) or SuperSignal West Femto (Thermo Scientific, Cat#34095) substrates. Images were acquired using a ChemiDoc MP imaging system (Bio-Rad).

### Small Molecule Inhibition

DIvA AsiSI-ER-U2OS cells were treated with various small molecule inhibitors, with or without the addition of 300 nM 4-Hydroxytamoxifen (4-OHT) to the cell culture media to induce breaks. Typically, cells were treated for 4H for ChIP and for 24H for QIBC, unless otherwise specified. For p53 inhibition, Pifithrin-α (506132-5MG) was used at 20 and 30 µM, for 24 hours.

### Cas9 and gRNA Preparation

Cas9-NLS (Streptococcus pyogenes) was obtained from QB3 MacroLab (UC Berkeley). Guide RNAs (gRNAs) were synthesized by annealing and PCR-amplifying single-stranded DNA oligonucleotides to generate a DNA template containing a T7 promoter, a 20 bp target-specific protospacer, an 80 bp scaffold, and a poly(A) tail. The template was transcribed in vitro using T7 RNA polymerase (HiScribe T7 Quick High Yield RNA Synthesis Kit; New England Biolabs, Cat#E2050S), followed by DNase I treatment to remove residual DNA. The 5′ triphosphate groups were removed by incubation with Shrimp Alkaline Phosphatase (New England Biolabs, Cat#M0371S), and gRNAs were purified using the RNeasy Mini Kit (QIAGEN, Cat#74106) for small-scale production or the Total RNA Purification Maxi Kit (Norgen Biotek, Cat#26800) for large-scale production.

### Cas9 RNP Assembly and Nucleofection

For recombination assays, 300 pmol of in vitro–transcribed SceGFP gRNA and 60 pmol of purified Cas9-NLS (5:1 molar ratio; QB3 MacroLab) were incubated for ≥10 min in 1× RNP buffer (20 mM HEPES pH 7.5, 200 mM KCl, 5 mM MgCl₂, 5% glycerol, 1 mM TCEP) in a final volume of 5 μL. U2OS DR-GFP cells (∼2.5 × 10 ) were harvested, pelleted (500 × g, 3 min), washed with D-PBS, and resuspended in 15 μL Nucleofection Buffer SE (Lonza, Cat#V4SC-1096). The RNP and cell suspensions were combined (total 20 μL) and electroporated using program CM-104 on a 4D-Nucleofector X Unit (Lonza). Post-nucleofection, cells were transferred to 6-well plates with pre-warmed recovery medium. Recombination rates were assessed 96 h post-electroporation by flow cytometry (Attune NxT, Invitrogen).

### Flow Cytometry

Flow cytometric analysis was performed on an Attune NxT Flow Cytometer (Invitrogen) and analyzed using FlowJo v10.7.1. Cells were gated for viability using forward scatter (FSC-A) versus side scatter (SSC-A), and singlets were identified using SSC-H versus SSC-A. For recombination assays in U2OS DR-GFP cells, GFP-positive cells were quantified from the singlet population using FSC-A versus BL1-H.

### Immunofluorescence Microscopy and Cell Cycle Analysis by QIBC

Cells were cultured in glass-bottom 96-well plates (Cellvis, Cat#P96-1.5H-N) to 50-80% confluency at the time of fixation and pulsed with 10 μM EdU for 30 min prior to fixation. Where indicated, pre-extraction was performed with ice-cold CSK buffer (10 mM PIPES–KOH pH 6.8, 100 mM NaCl, 10% sucrose, 1 mM EGTA, 1 mM MgCl₂, 0.25% Triton X-100) for 2 min. Cells were fixed in 4% formaldehyde in D-PBS for 10 min, permeabilized with 0.25% Triton X-100 in D-PBS for 10 min, and subjected to click chemistry in RB buffer (50 mM Tris–HCl pH 7.5, 150 mM NaCl) with 2 mM copper sulfate, 5 μM azide, and 1 mg/mL sodium ascorbate for 30 min at room temperature in the dark. Following washes, cells were blocked in 3% BSA/0.1% Tween-20 in D-PBS and incubated with primary antibodies overnight at 4°C, followed by secondary antibodies and DAPI for 1 h at room temperature. Images were acquired on a Nikon Ti2-E inverted microscope with a Yokogawa CSU-W1 spinning disk unit and ORCA-Fusion BT sCMOS camera using a 10× plan apo Lambda D objective (NA 0.45) for QIBC or a 100× oil immersion objective (NA 1.49) for high-magnification imaging. Only the central quarter of each image was retained to avoid vignetting, and 9–16 images were collected per well.

QIBC analysis was performed in CellProfiler (v4.2+ Broad Institute), with background subtraction based on the 10% intensity value, nuclear segmentation from DAPI staining, and foci enhancement (feature size 5) prior to thresholding. Quantification and visualization were performed in Python v3.12 using Pandas v1.5.3 and Seaborn v0.13.2.

### Chromatin Immunoprecipitation

Cells (5 × 10 ) were crosslinked in serum-free medium with 1% formaldehyde (Thermo Scientific, Cat#28908) for 10 min at room temperature, quenched with 0.125 M glycine for 5 min, and washed three times with ice-cold D-PBS. Pellets were either processed immediately or flash frozen at −80°C.

For lysis, pellets were resuspended in 1 mL LB1 (50 mM HEPES–KOH pH 7.5, 140 mM NaCl, 1 mM EDTA, 10% glycerol, 0.5% IGEPAL CA-630, 0.25% Triton X-100) with protease inhibitors (Fisher Scientific, Cat#PI78429) and incubated for 10 min at 4°C with rotation. Cells were centrifuged (2000 × g, 3 min, 4°C), and pellets were resuspended in 1 mL LB2 (10 mM Tris–HCl pH 8.0, 200 mM NaCl, 1 mM EDTA, 0.5 mM EGTA) with protease inhibitors, incubated for 5 min at 4°C with rotation, and centrifuged again (2000 × g, 3 min, 4°C). Pellets were then resuspended in 0.5 mL LB3 (10 mM Tris–HCl pH 8.0, 100 mM NaCl, 1 mM EDTA, 0.5 mM EGTA, 0.1% sodium deoxycholate, 0.5% N-lauroylsarcosine) with protease inhibitors.

Chromatin was sheared in LB3 using a Qsonica Cup Horn Sonicator (15 s on/45 s off, total on-time 6 min, 25–50% amplitude) to achieve 200–600 bp fragments. 25 μL of each sonicated sample was collected for a sonication size check. Lysates were clarified by centrifugation (21,100 × g, 10 min, 4°C), adjusted to 1 mL with LB3 containing 1% Triton X-100, and 5% was reserved as input. Immunoprecipitations were performed overnight at 4°C with 50 μL Dynabeads Protein A or G (Invitrogen) pre-bound to 5–10 μg antibody.

Cleared lysates were adjusted to 1 mL with LB3 containing 1% Triton X-100, and 5% was reserved as input. Immunoprecipitations were performed overnight at 4°C with 50 μL Dynabeads Protein A or G (Invitrogen) pre-bound to 5–10 μg of antibody. Beads were washed five times with ChIP wash buffer (50 mM HEPES–KOH pH 7.5, 500 mM LiCl, 1 mM EDTA, 1% IGEPAL CA-630, 0.7% sodium deoxycholate) and once with TBS (20 mM Tris–HCl pH 8.0, 150 mM NaCl).

The following day, immunoprecipitated samples were washed five times with ice-cold ChIP wash buffer (50 mM HEPES–KOH, pH 7.5; 500 mM LiCl; 1 mM EDTA; 1% IGEPAL CA-630; 0.7% sodium deoxycholate) followed by a single wash with ice-cold TBS (20 mM Tris–HCl, pH 8.0; 150 mM NaCl). Bound chromatin was then eluted twice at 65°C with shaking at 1200 rpm for 15 min per elution using 60 μL of ChIP elution buffer (0.1 M NaHCO₃, 1% SDS) for each round. To reverse protein–DNA crosslinks, the combined eluates were incubated overnight at 65°C with shaking (1200 rpm) in the presence of 200 mM NaCl and 2 μL RNase A (10 mg/mL). The following day, 2 μL Proteinase K (20 mg/mL) was added, and samples were incubated for 1 h at 65°C with shaking (1200 rpm). DNA was then purified using the MinElute PCR Purification Kit (QIAGEN, Cat#28004) and analyzed for enrichment by qPCR.

### qPCR

Quantitative PCR was used to assess ChIP enrichment. Each reaction contained 3 μL of ChIP DNA, 1.9 μL nuclease-free water, 0.05 μL each of 100 μM forward and reverse primers, and 5 μL of 2× SsoAdvanced Universal SYBR Green Supermix (Bio-Rad, Cat#1725271). Reactions were run for 45 cycles using a two-step protocol, followed by melting curve analysis from 65°C to 95°C in 0.5°C increments with 5 s per step.

Samples with amplification detected after cycle 35 were considered negative.

### Stranded Library Prep

For selected ChIP samples, stranded DNA libraries were prepared using the xGen ssDNA & Low-Input DNA Library Prep Kit, 96-reaction format (Integrated DNA Technologies, Cat#10009817) following the manufacturer’s instructions.

### mRNA-seq

DIvA U2OS cells were transfected with 20 pmol siRNA targeting SFPQ or a non-targeting control (see RNA interference section). DNA breaks were induced with 4-OHT or left uninduced. Total RNA was extracted from ∼1 × 10 cells using the Direct-zol RNA Miniprep Kit (Zymo Research, Cat#R2050) following the manufacturer’s protocol, and eluted RNA was stored at −80°C. Samples were submitted to Novogene for human mRNA sequencing, including poly(A) enrichment and library preparation, and sequenced on a NovaSeq X Plus platform (PE150) to a depth of 6 G raw reads per sample.

### ChIP-seq data analysis

Reads were quality and adaptor trimmed using fastp (v0.23.2), aligned to the hg38 genome using bowtie2 (v2.5.3), and deduplicated using picard. The resulting bam files were processed in experiment-level batches using the deepTools (v3.5.1) packages multiBamSummary --scalingFactors and bamCoverage --scaleFactor to produce TMM normalized bigwig files.

Signal at multiple genomic loci in each TMM normalized bigwig file was combined using the deepTools computeMatrix and plotProfile commands. BED files specifying regions of interest are presented in the Supplemental Table. Features with approximately equal sizes (DSBs, RNUs) were processed using the reference-point subcommand of computeMatrix. Features with variable sizes (RAD51 paralogs) were processed with the scale-regions subcommand of computeMatrix, which fits signal from features with different lengths into the same plot width.

### RNAseq data analysis

Reads were quality and adaptor trimmed using fastp (v0.23.2), aligned to the hg38 genome using STAR (v2.7.11a) quantmode GeneCounts, and reads per gene count tables were combined into a single count table file. Differential expression was calculated using the DESeq2 R package (v. 1.48.2) and plotted using Seaborn.

### Differential transcript utilization

RNAseq reads were analyzed using a custom shell script that aligned reads to the gencode.v48 transcriptome using Salmon (v.1.2) and evaluated differential transcript utilization using DRIMseq (v1.36). Differential transcript utilization was calculated at the gene level and likelihood ratios and adjusted p-values were plotted using Seaborn.

### RIPseq data analysis

Reads (SRR14127799, SRR14127800, SRR14127801, SRR14127802) were quality and adaptor trimmed using fastp (v0.23.2) and aligned to the hg38 genome using bowtie2 (v2.5.3). Reads mapping to the ENCODE blacklist sites (hg38-blacklist.v2.bed) were removed. The resulting bam files were displayed using IGV and exported as SVG images.

### Data Availability

Raw data for ChIP and RNAseq have been uploaded to SRA under SUB15601024 and will be available upon publication.

Raw data for RIP-seq was reprocessed from PRJNA719095.

## Acknowledgements

We are grateful to Gaëlle Legube (Center for Integrative Biology, Toulouse, France) for sharing DIvA AsiSI-ER-U2OS cells, and to Jeremy Stark (City of Hope, Duarte, California, USA) for providing U2OS DR-GFP cells. We also thank the Gardner and Richardson laboratory members for their valuable feedback on this work. We further acknowledge Dr. Jennifer Smith, manager of the Biological Nanostructures Laboratory at the California NanoSystems Institute, which is supported by the University of California, Santa Barbara, and the University of California, Office of the President.

## Author Contributions

**S. Gotthold:** Conceptualization, Investigation, Validation, Visualization, Data Curation, Writing - Original Draft, Writing - Review & Editing. **K. R. Hansen:** Investigation, Conceptualization, Validation, Writing - Review & Editing. **A. N. Brown:** Investigation, Formal analysis, Software, Validation, Writing - Review & Editing. **S. P. Chowdhury:** Investigation, Conceptualization, Formal analysis, Writing - Review & Editing. **H. I. Ghasemi:** Investigation, Writing - Review & Editing. **A. C. Yoon:** Investigation, **C. M. Joyce:** Investigation, Writing - Review & Editing. **J. Bacal:** Investigation, Conceptualization, Formal analysis, Software, Writing - Review & Editing. **B. M. Gardner:** Conceptualization, Methodology, Funding acquisition, Project administration, Supervision. **C. D. Richardson:** Conceptualization, Methodology, Software, Validation, Investigation, Formal analysis, Funding acquisition, Project administration, Supervision, Visualization, Data Curation, Resources, Writing - Review & Editing.

